# Transcription factors drive opposite relationships between gene age and tissue specificity in male and female *Drosophila* gonads

**DOI:** 10.1101/2020.12.09.418293

**Authors:** Evan Witt, Nicolas Svetec, Sigi Benjamin, Li Zhao

## Abstract

Evolutionarily young genes are usually preferentially expressed in the testis across species. While it is known that older genes are generally more broadly expressed than younger genes, the properties that shaped this pattern are unknown. Older genes may gain expression across other tissues uniformly, or faster in certain tissues than others. Using *Drosophila* gene expression data, we confirmed previous findings that younger genes are disproportionately testis-biased and older genes are disproportionately ovary-biased. We found that the relationship between gene age and expression is stronger in the ovary than any other tissue, and weakest in testis. We performed ATAC-seq on *Drosophila* testis and found that while genes of all ages are more likely to have open promoter chromatin in testis than in ovary, promoter chromatin alone does not explain the ovary-bias of older genes. Instead, we found that upstream transcription factor (TF) expression is highly predictive of gene expression in ovary, but not in testis. In ovary, TF expression is more predictive of gene expression than open promoter chromatin, whereas testis gene expression is similarly influenced by both TF expression and open promoter chromatin. We propose that the testis is uniquely able to expresses younger genes controlled by relatively few TFs, while older genes with more TF partners are broadly expressed with peak expression most likely in ovary. The testis allows widespread baseline expression that is relatively unresponsive to regulatory changes, whereas the ovary transcriptome is more responsive to trans-regulation and has a higher ceiling for gene expression.

## Introduction

For eons, genes have continuously arisen by a multitude of ways, from duplication and divergence to *de novo* origination from non-genic DNA (Begun et al., 2006; Long et al., 2003; Ohno, 1970; Tautz and Domazet-Lošo, 2011; Zhao et al., 2014; Zhou et al., 2008). Gene birth and death is a continuous and dynamic process in evolution, culled by natural selection or genetic drift (Kaessmann, 2010; Palmieri et al., 2014). A large portion of young genes segregate within or recently fixed in populations, and most young genes are expressed specifically in the testis (Levine et al., 2006; Zhao et al., 2014), similar to duplicated genes (Long et al., 2013). The phrase “out of the testis” was originally used to describe young retroposed genes (Vinckenbosch et al., 2006), which gained expression by exploiting cis-regulatory machinery of nearby genes. Testis-bias has since been observed in young X-linked duplicate genes, leading researchers to propose that young genes escape Meiotic Sex Chromosome Inactivation (MSCI) due to immature cis-regulatory machinery (Zhang et al., 2010a). Testis expresses more genes in general than any other tissue (Soumillon et al., 2013), and studies from many taxa support that a large proportion of young genes then to show testis-biased or testis-specific expression and function (see review in Long et al., 2013).

The testis-biased expression of young genes has many possible explanations. Besides the obvious hypothesis that genes expressed in reproductive tissues may directly influence reproductive success and fitness (Begun et al., 2006; Zhang et al., 2004), many propose that the testis has a permissive chromatin environment facilitating the transcriptional birth of genetic novelties (Kaessmann, 2010; Soumillon et al., 2013). Indeed, most genes are at least somewhat expressed in the testis (Soumillon et al., 2013; Witt et al., 2019). It has long been proposed that an upregulation of universal transcriptional machinery facilitates such widespread transcription (Schmidt, 1996). Such broad transcription may be a form of genomic surveillance, meant to detect and repair mutations via transcription-coupled repair or other mechanisms (Grive et al., 2019; Xia et al., 2020). It has also been proposed that permissive testis transcription is also due to reduced mRNA degradation of testis-specific genes (Mayr, 2016). Young genes may also have low levels of “active” epigenetic markers across tissues, despite high expression in testis (Zhang and Zhou, 2019). Results from Zhang and Zhou 2019 suggest that young genes have similar epigenetic profiles across tissues, yet show testis-biased expression, while older genes show consistently higher levels of “active” epigenetic marks. Their results indicate that the “out of the testis” pattern for the emergence of young genes may not be driven by specific epigenetic marks, but rather by a context-dependent trans-regulatory environment between tissues (Ding et al., 2010). Alternatively, recruitment of nearby testis-biased cis-regulatory elements by young genes may also be responsible for many testis-biased new genes (Majic and Payne, 2020).

While it is known that young genes are often testis-specific, and that older genes are more broadly-expressed than young genes (Kondo et al., 2017; Zhou et al., 2008), it is unknown how this relationship works. When genes age, do older genes lose expression in testis, and retain relatively constant expression in other tissues? Or do older genes maintain relatively constant expression in testis, and gain expression in other tissues? If so, are all non-testis tissues equally conducive to old genes, or do the genomic characteristics of older genes produce higher expression in certain tissues? Once out of the testis, is any tissue the next hot target of tissue-biased expression when the genes expand their functions in other tissues?

One clue is that older duplicated genes are more likely to be retained if they are ovary-biased (Assis, 2019). This might imply a specific importance of older genes to ovary expression and function. To this effect, researchers have identified several modules of highly conserved, older genes with heightened importance in human ovarian function (Zhang et al., 2019). To see if the *Drosophila* ovary drives the shift away from testis-bias in older genes, we analyzed a database of RNA-seq data from FlyAtlas2 (Leader et al., 2018) to characterize tissue-bias for genes of all ages in every tissue. We found that ovary has the largest relationship between gene age and expression, explaining why the oldest genes are often ovary biased. Conversely, testis shows a weaker relationship between gene age and gene expression than any other tissue.

To explain this trend, we examined the tissue-specific activity of the transcription factor (TF) regulators of every gene in the DroID database (Murali et al., 2011). We found that ovary-biased genes tend to have higher upstream TF expression than testis-biased genes of all age groups, yet young genes, with fewer TF partners, tend to be testis-expressed and old genes, with more TF partners, tend to be ovary biased. We found evidence that testis allows higher transcription than the ovary for genes with low TF expression. Conversely, genes with high TF expression have higher expression in the ovary than the testis. Additional upstream TF expression appears to confer diminishing returns on expression in testis, but greatly benefits ovary expression, explaining why older genes with more TF partners tend to be ovary biased.

After establishing the different relationships between trans regulation and gene expression in testis and ovary, we performed ATAC-seq to assess if open promoter chromatin is equally predictive of expression in the two tissues. All age groups of genes are more likely to have open promoter chromatin in testis than ovary, indicating that open chromatin by itself is insufficient to explain age-related expression bias. In ovary, we found that high upstream TF expression is much more predictive of gene expression than the presence of open promoter chromatin, whereas in testis, high TF expression and open promoter chromatin are similarly predictive of gene expression. This indicates that gene expression in ovary is much more linked to trans-regulatory factors than testis expression. Taken together with our observation that young genes are less likely to be bound by annotated TFs than older genes, the opposite trends of gene age and tissue bias in testis and ovary make biological sense. We published a web app to allow users to interactively explore our tissue specificity data for any set of genes without coding experience necessary: https://zhao.labapps.rockefeller.edu/tissue-specificity/.

## Results

### Testis and ovary show an opposite relationship between gene age and tissue bias

Using gene ages divided into Drosophilid (youngest), pre-Drosophilid (middle-aged), and pre-Bilateria (oldest), and tissue RNA-seq data from FlyAtlas2, we find results consistent with earlier work showing that younger genes are more tissue specific than older genes (Figure 1A). We plotted the proportion of genes from each age group with maximum expression in testis, ovary, and male and female carcasses with the reproductive tracts removed. A plurality of young genes are testis biased, but the abundance of testis-biased genes declines for older genes (Figure 1B). Surprisingly, we found the opposite trend for ovary: older genes are very likely to have maximum expression in ovary, but almost no younger genes are ovary biased. No other tissues displayed a relationship of this magnitude (Supplemental Figure 1), indicating that the two tissues that contribute most to gene age-related expression patterns are the male and female reproductive tissues. Whereas young genes are often testis-biased and highly tissue specific, old genes are broadly expressed with peak expression in ovary.

**Figure 1:**
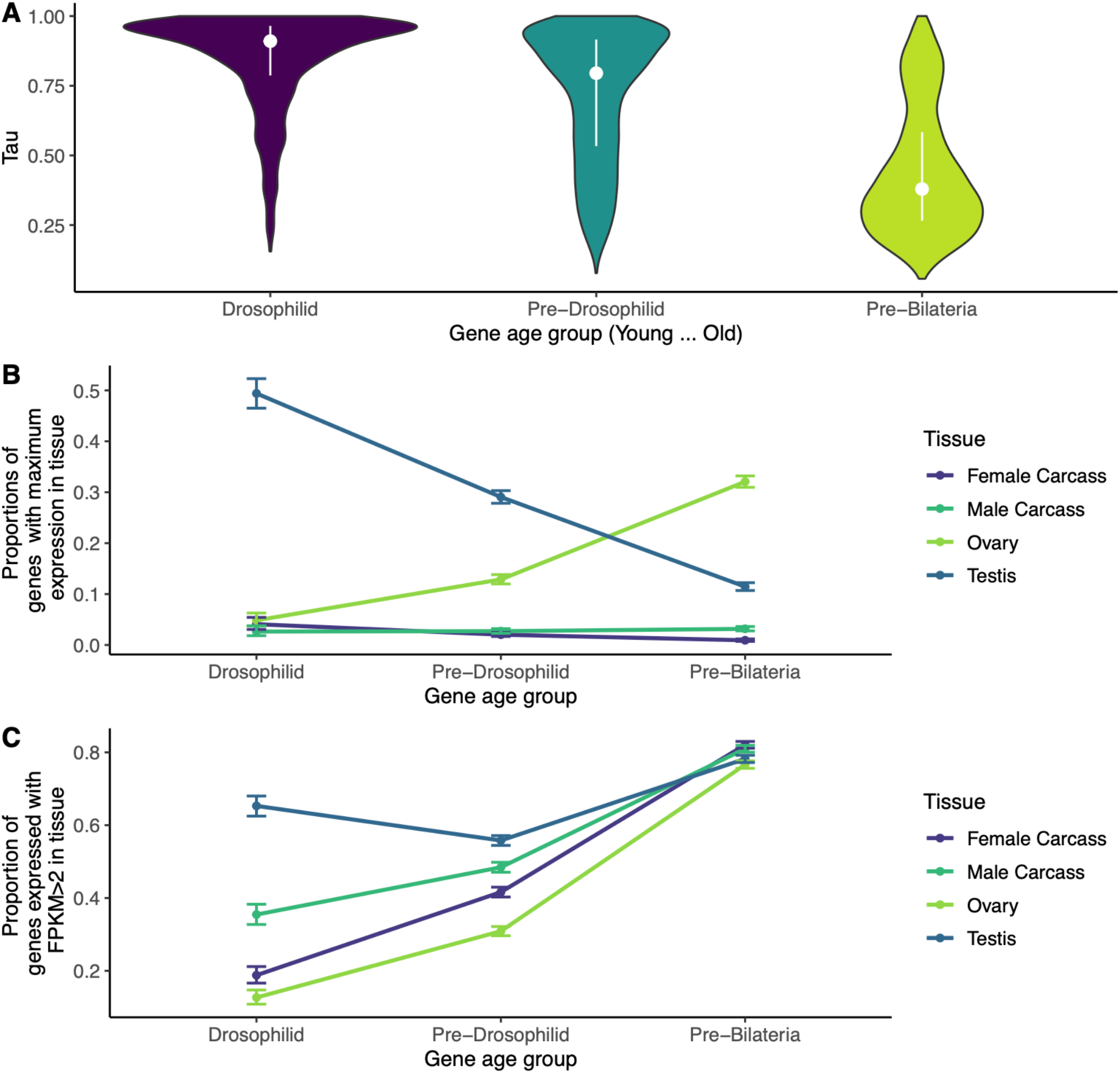
Young genes are testis-specific, old genes are broadly-expressed and often ovary-biased. A) Average Tau values among genes of each age group, dots represent medians and vertical lines are interquartile ranges. Young genes are more tissue specific (higher Tau) than older genes. B) For four tissues, proportion of genes of each age group with maximum expression in that tissue. Younger genes usually have highest expression in testis, but the proportion of testis-biased genes declines with gene age. Ovary biased young genes are rare, but old genes are more often biased towards ovary than any other tissue. Error bars are 95 percent confidence intervals for proportion test. C) For four tissues, proportion of genes of each age group with FPKM >2 in that tissue. In testis, ovary and carcass, old genes are more likely to be expressed than young genes, but this disparity is smallest in testis and largest in ovary. By this measure, old genes are no longer biased in testis. Ovary-bias of older genes is not explained by the relative proportion of genes expressed between tissues.

While a plurality of old genes are ovary-biased, this is not due to an increased likelihood of expression for old genes in ovary. Young genes are most commonly expressed with FPKM>2 in testis (65%) and least commonly expressed in ovary (13%), whereas testis, ovary and somatic tissues express a similar proportion of old genes (all between 73% and 85%; Figure 1C, Supplemental Figure 2). Therefore, the age-related decline in testis-bias is not due to an absence of old gene expression in testis. The proportion of genes expressed between age groups varies the least in testis, and the most in ovary, indicating that ovary may have a disproportionately large relationship between gene age and expression. We confirmed that young duplicate genes were not confounding these results by repeating the analysis from Figure 1 with *melanogaster*-specific genes removed (Supplemental Figure 3). We also confirmed these results with an alternate set of gene age assignments (Supplemental Figure 4).

### Testis shows a weak, and ovary shows a strong relationship between gene age and expression

We wanted to further unpack how gene expression correlates with gene age across tissues to understand our observed patterns of testis-bias and ovary-bias. For each tissue, we plotted gene expression (Log2(FPKM +1)) from FlyAtlas2 conditioning by gene age. In every tissue, expression of old genes was higher for pre-Bilateria genes than for Drosophilid genes as measured with a pairwise Wilcoxon test (Figure 2A). In every tissue except testis, Drosophilid genes were less expressed on average than pre-Drosophilid genes. In testis, these two groups were statistically similar. This may be because in testis, unlike other tissues, a similar proportion of genes are expressed for each age group (Figure 1C). A qualitative comparison shows that expression of the 3 age groups is least different in testis (median FPKM 8.53 (Drosophilid), 3.19 (Pre-Drosophilid), 8.24 (pre-Bilateria)), and most dramatically different in ovary (median FPKM 0.071, 0.25, 21.10 respectively) (Figure 2A).

**Figure 2:**
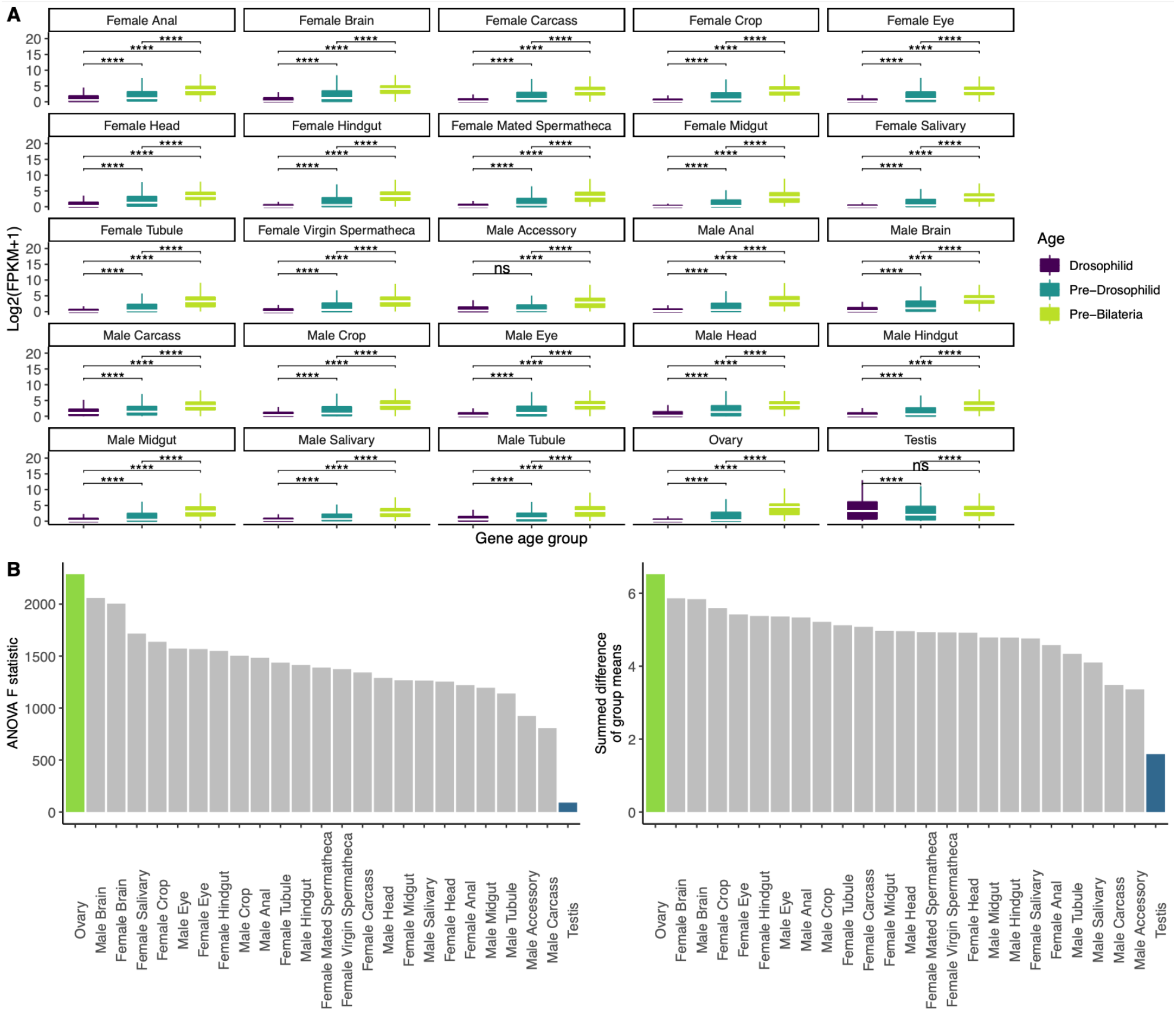
Tissue-specific trends in the gene-age/expression relationship. A) The log-scaled expression of every gene in that tissue versus gene age for every adult tissue (see methods). In almost every tissue scaled expression is very low for young genes, and high for older genes. The testis is an outlier, with statistically similar expression between Drosophilid and Pre-Drosophilid genes. Asterisks represent P values are adjusted with Bonferroni’s correction (*=p<0.05, **=p<0.005,***=p<0.0005, ****=p<0.00005). Raw and Bonferroni-adjusted p values are in Supplemental Table 1. B) Rankings of ANOVA statistics for all tissues. We performed an ANOVA on each of the panels from part A, comparing, for each tissue, the ratio of inter-group variation to between group variation (F statistic). By this measure, ovary has the largest relationship between gene age and expression (because old genes are often ovary biased), and testis has the smallest (because old and young genes are similarly expressed in the testis). We also took the mean difference between groups and summed their absolute values for each tissue. Testis has the smallest mean expression difference between age groups, and ovary has the largest. This conclusion held when we repeated the analysis using an alternate set of gene age assignments (Supplemental Figure 6).

To quantitatively compare tissue-specific gene expression as a function of age group, we performed a one-way ANOVA on each tissue and age group from Figure 2A. The ANOVA F statistic is the ratio of between group variation to intra-group variation. For similar groups, the F statistic is close to 1. The ANOVA F statistic is highest in ovary, meaning that the age groups are more variable in this tissue than any other. In testis, the F statistic is lower, meaning that gene expression varies less between age groups. Young genes have relatively similar expression in testis across all age groups, in contrast to other tissues where gene expression is highly stratified across age groups, with young genes the least and old genes the most expressed.

For each tissue, we also calculated the summed pairwise mean differences between every group. This measure is the absolute value of the difference between the mean of each age group within a tissue, summed for each pair of groups (Drosophilid vs. Pre-Drosophilid, Pre-Drosophilid vs. Bilateria, Drosophilid vs. Bilateria). By this measure, mean testis expression is the least different between gene age groups and ovary expression varies the most of any tissue (Figure 2B). The results in Figure 2B hold if *melanogaster-*specific genes are removed (Supplemental Figure 5), or with an alternate method of gene age assignments (Supplemental Figure 6).

### Testis expression requires lower transcription factor activity than ovary expression

We hypothesized that TFs may play a role in the discrepancy between age/expression relationships between the testis and ovary. We designed a proxy measure of TF network activity for every gene in every tissue. For every gene with bound TFs listed by DroID (Murali et al., 2011), we defined the summed scaled expression of the upsteam TFs of a gene in a tissue as “TF expression”. Higher TF expression in a tissue indicates that a gene’s TF partners are more transcriptionally active in that tissue. This metric is based on data from ChIP-seq and ChIP-chip experiments for individual TFs and only considers whether a TF binds to a given gene’s promoter. While such a method does not reveal whether a TF-gene relationship is one of activation or repression, it is unbiased with regard to gene age since the whole-genome binding profile of a TF is agnostic to the degree of study a particular gene has received (as young genes are often less studied than older genes with mammalian homologs).

The purpose of our TF expression metric is not to infer gene expression (for which RNA-seq is much better suited), but rather to assess the relative dynamics between gene expression and trans regulation across tissues. For this purpose, the metric performs consistently well across tissues even though some TFs are repressive in nature. For more details about TF expression, see methods.

We compared the TF expression of young, middle-aged, and older genes between the testis and ovary. We thought that since young genes are more specifically expressed in testis, young genes would have higher uptsream TF expression in testis than in ovary. We found that no age group of genes shows higher TF expression in testis than ovary (Figure 3A). The testis-specificity of young genes must be due to factors other than increased TF expression in testis. Exploring further, we found that young genes have fewer identified TF-gene interactions than middle-aged genes, which in turn have fewer TF binding partners than old genes (Figure 3B). We confirmed these results using an alternate list of gene ages in Supplemental Figure 7.

**Figure 3:**
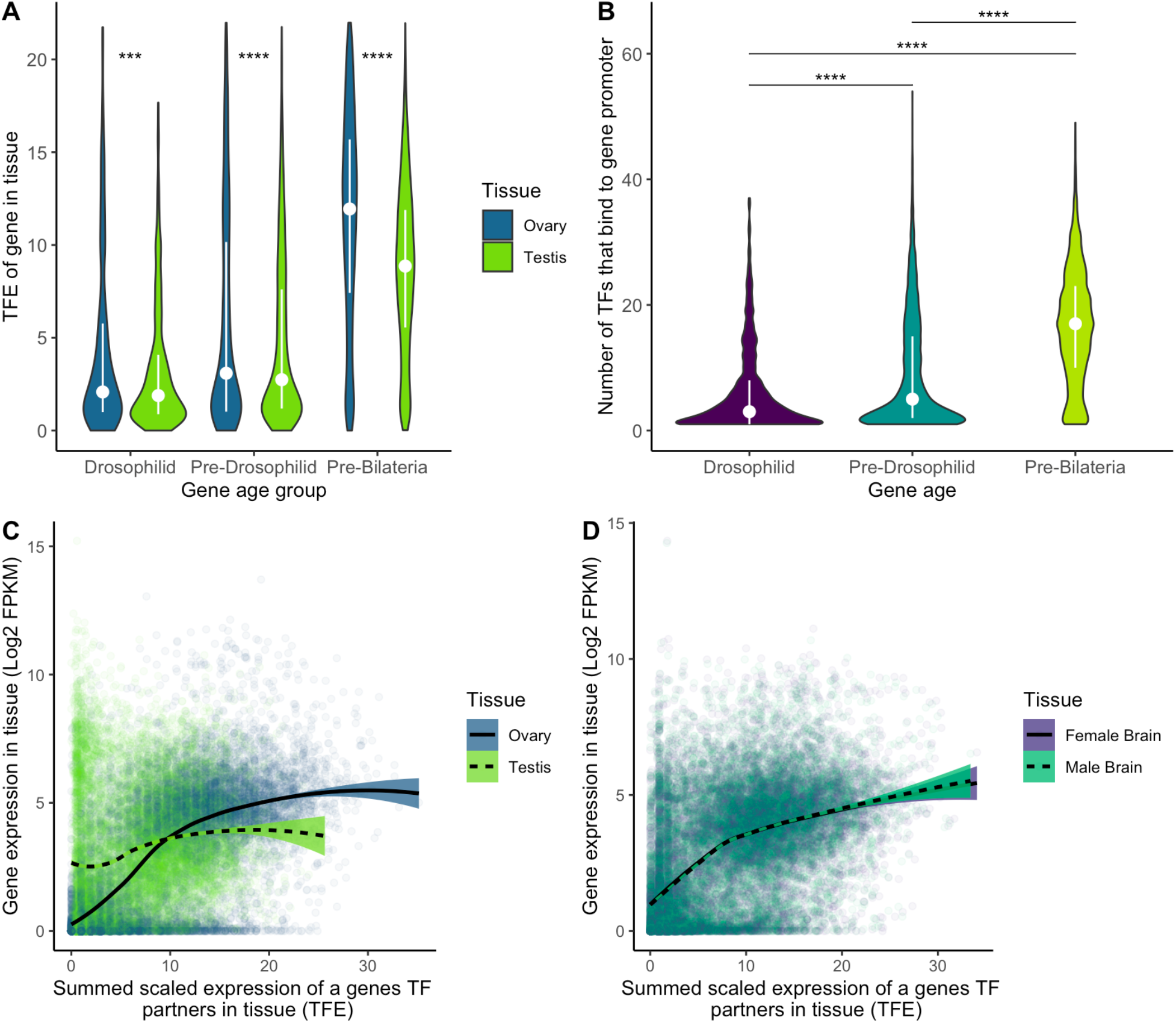
Transcription factor expression explains gene age/expression trends in testis and ovary. A) Upstream TF expression in testis and ovary for genes with different ages, using DroID’s curated database of TF binding profiles from the modENCODE project. For every gene with a confirmed TF-promoter interaction, we calculated TF expression in testis and ovary by scaling the expression for each TF from 0 to 1, and summing the scaled expression of every TF that binds to the promoter of a given gene. Older genes have much higher TF expression than younger genes in both tissues, and no age group of genes shows elevated TF expression in testis compared to ovary. White dots are medians and lines are interquartile ranges. Asterisks represent p values adjusted with Bonferroni’s correction. B) In DroID data, the promoters of older genes have been shown able to be bound by more TFs than younger genes. C) Log-scaled gene expression versus upstream TF expression in gonads. Genes require less upstream TF expression for expression in testis than in ovary. Genes have fairly high testis expression even without much TF expression in testis, but genes with low ovary TF expression are relatively lowly expressed in ovary. This indicates that genes require less TF expression for testis expression than ovary expression. In ovary, higher TF expression corresponds to higher expression, moreso than testis, where adding TF expression makes relatively little difference in testis gene expression. D) Log-scaled gene expression versus upstream TF expression in brains. These sexually dimorphic tissues show no difference in their relationships between TF expression and gene expression. In these tissues, low TF expression yields low expression, and high TF expression yields high expression, much like the ovary and much unlike the testis. Lines are smoothed loess regressions with 95 percent confidence intervals. Other tissues are shown in Supplemental Figure 3.

We then sought to correlate expression with TF activity between testis and ovary, and found that genes with low TF expression are much more active in testis than in ovary. Conversely, genes with high TF expression are often more active in the ovary than in testis (Figure 3C). It appears that testis expression requires fewer TFs than ovary expression, explaining why young genes, with fewer TFs, would have testis-biased expression. Having many TF partners, a property of older genes, appears to boost expression in ovary more than in testis. To confirm that this property was not sex-specific we compared TF expression and gene expression in the male and female brain, two sexually dimorphic tissues, and observed no major differences (Figure 3D). Additionally, we made this comparison across all tissues in FlyAtlas2 (Supplemental Figure 8), and found that gene expression is least correlated to TF expression in testis (Pearson’s r=0.22), and most responsive in ovary (Pearson’s r=0.67).

### Testis promoter chromatin is broadly open across all gene ages

To see whether promoter chromatin environment explains TF expression differences in testis and ovary, we performed ATAC-seq on *Drosophila* testis, and obtained ATAC-seq datasets for *Drosophila* Ovarian Somatic Cells (Iwasaki et al., 2016), and S2 cells (Vaid et al., 2020). We annotated peaks in the promoters of genes from each age group, and compared the proportion of genes with detectable high-quality peaks in each tissue (Figure 4A). In every tissue, young genes were the least likely to have detectable chromatin accesibility in their promoters, and old genes were the most likely to have detectable peaks. Every age group of genes was more likely to have peaks in testis, and least likely to have peaks in ovary, indicating that chromatin at the promoter is more broadly open in the testis. In addition, a majority of genes from each age group exhibited more frequent detectable open promoter chromatin in testis. In ovary, by contrast, pre-Bilateria genes are the only age group of which a majority of genes (68%) have detectable ATAC-seq peaks. Every other age group of genes is less likely to contain detectable promoter ATAC-seq peaks, especially young genes, of which only 26% have open chromatin in ovary, compared to 56% of young genes in testis. Our observation that every gene-age group is more likely to have testis peaks than ovary peaks indicates that open chromatin does not underlie the ovary-bias of older genes.

**Figure 4:**
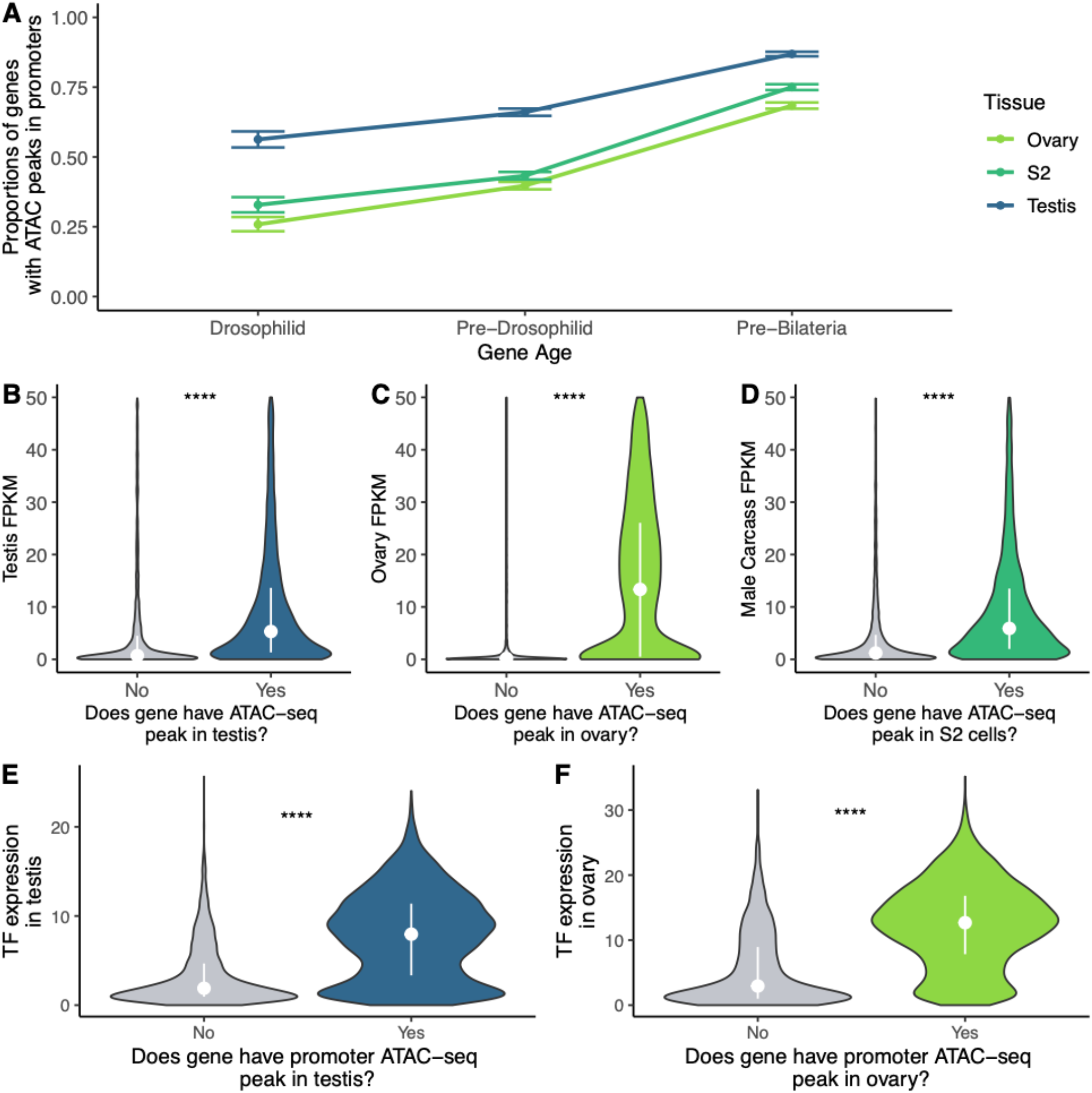
ATAC-seq peaks show an age-related trend in multiple tissues. A) The relative proportions of genes with a detectable ATAC-seq peak in their promoters, for 3 gene age groups and 3 datasets. For each dataset, young genes were the least likely to have open chromatin in their promoters. Testis is unique among these datasets because a majority of genes of each age group have open promoter chromatin. B) FPKM for genes with and without a detected promoter ATAC-seq peak in testis. Genes with open promototer chromatin in testis have generally higher expression in FlyAtlas2 data. Dots are medians, white line is the interquartile range. C) Genes with open promoter chromatin in ovary have higher FlyAtlas2 expression, and the FPKM difference between genes with and without peaks is much larger than the other two tissues. D) Genes with promoter peaks in S2 cells generally have higher expression in male carcass, the most analogous FlyAtlas2 tissue to this cell line. E) TF expression for genes with and without detectable ATAC-seq peaks. Genes with ATAC-seq promoter peaks tend to have higher TF expression in testis, F) as well as ovary. **** represents adjusted p values <0.00005.

The presence of an ATAC-seq peak generally corresponds to increased gene expression in analogous tissues (Figure 4B, 4C, and 4D). Similarly, genes with an ATAC-seq peak in a tissue have heightened activity of their partner TFs compared to genes with no peak in a tissue (Figure 4E, 4F). This indicates that TF expression and promoter chromatin state are useful proxies of a gene’s network activity (Sigalova et al., 2020).

The low proportion of young genes with ovary ATAC-seq peaks does not entirely explain the paucity of young ovary-biased genes. In the ovary, we found that 13% of young genes are expressed while 26% of them have open promoter chromatin. We therefore sought next to separate the relative influences of TF expression and promoter chromatin for testis and ovary expression

### High upstream TF expression boosts gene expression in ovary more than in testis

We quantified expression for genes with and without detectable ATAC-seq peaks, conditioning on whether they had high or low TF expression in the tissue (Figure 5). Many genes in testis have surprisingly high expression (median FPKM 1.04) without nearby detectable ATAC-seq peaks or high TF expression, indicating that baseline transcription is higher in testis than in ovary (median FPKM 0.13). Without the aid of many TF partners or open promoter chromatin detectable by ATAC-seq, plenty of genes have surprisingly high expression in testis, but not ovary. In both testis and ovary, the presence of detectable ATAC-seq peaks or high TF expression (greater than the tissue median) is associated with an expression boost. In testis, however, these fold differences in median expression are smaller than in ovary (Table 1). Furthermore, in ovary, high TF expression boosts expression 169.23 fold in genes without a detectable ATAC-seq peak. For genes with a detectable ATAC-seq peak in ovary, high TF expression is associated with a further 15% boost in expression. Ovary expression is 24.18 fold higher for genes with high TF expression but no detectable ATAC-seq peaks compared to genes with open chromatin but low TF expression, indicating that high TF expression is more predictive of expression than chromatin environment in ovary.

**Figure 5:**
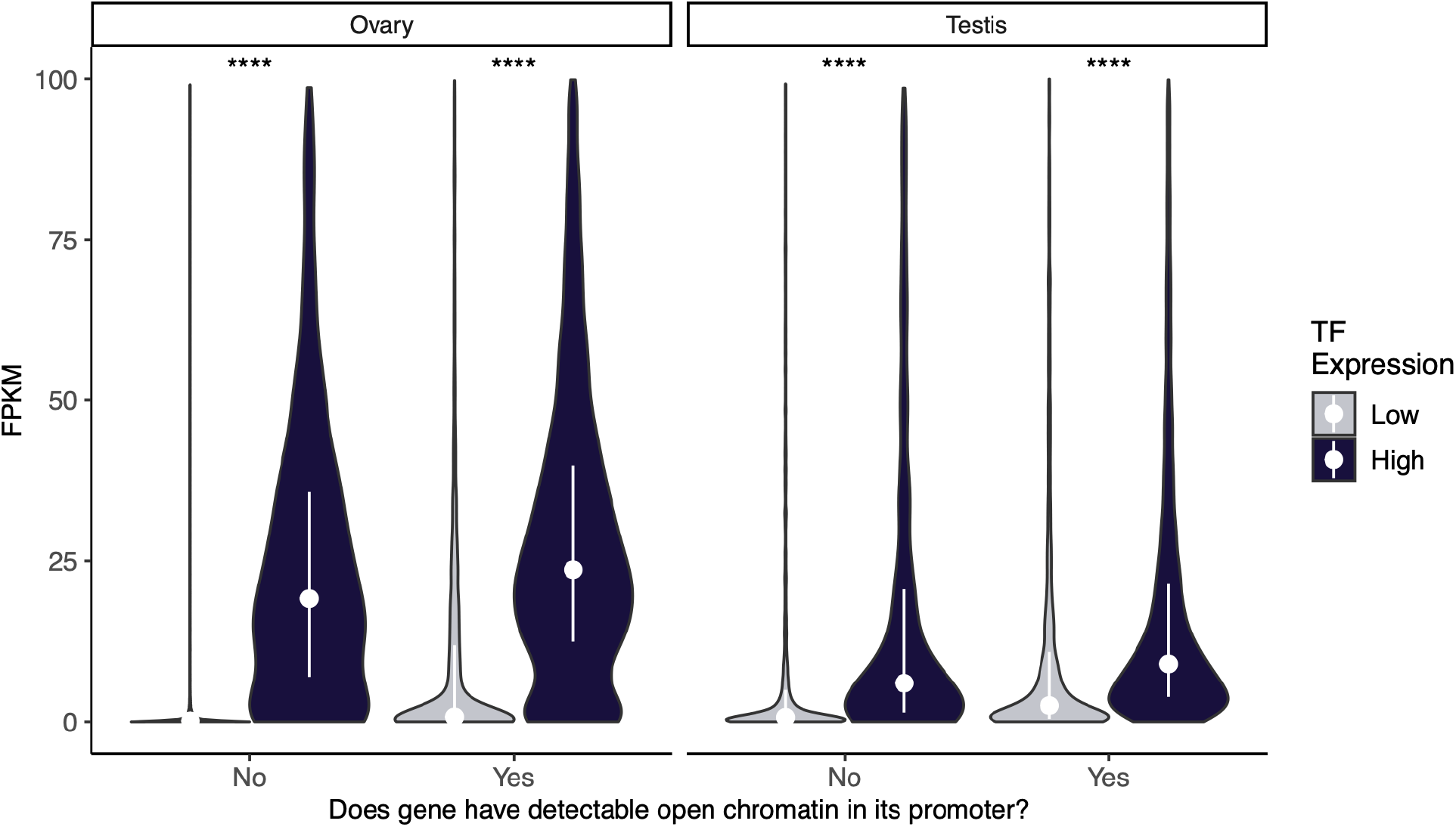
High TF expression disproportionately predicts gene expression in ovary. For testis and ovary, FPKM for genes with and without detectable chromatin peaks, grouped by “high” or “low” upstream TF expression. Genes are classified as high or low activity in a tissue if they are above or below the median TF expression for genes in the tissue. In testis, both high TF expression and open promoter chromatin confer a similar, modest expression benefit. In ovary, genes with low TF expression are generally very lowly expressed regardless of the presence of a promoter peak. This indicates that TF expression influences ovary expression more than chromatin environment. In ovary, high TF expression is necessary and sufficient for gene expression. White dots are the median values for each group, used to calculate fold changes in table 1. Vertical lines are inter-quartile ranges.

**Table 1:** High TF expression confers a disproportionate fold difference in gene expression in ovary. Corresponding to the median values shown in figure 5, these are the pairwise fold differences in median FPKM for genes with and without promoter peaks, and genes with upstream TF expression above or below the median for a tissue. In ovary, genes with no detectable peak have 169.23-fold higher expression if their TF expression is higher than the median TF expression for genes in ovary. In testis, fold differences in median expression are much smaller between groups.

| Category | Ovary median FPKM | Ovary fold difference | Testis median FPKM | Testis fold difference |
| --- | --- | --- | --- | --- |
| Low TF expression, no ATAC peak | 0.13 | 7.00 | 1.04 | 3.37 |
| Low TF expression, ATAC peak | 0.91 |  | 3.50 |  |
| Low TF expression, no ATAC peak | 0.13 | 169.23 | 1.04 | 6.92 |
| High TF expression, no ATAC peak | 22.00 |  | 7.20 |  |
| Low TF expression, no ATAC peak | 0.13 | 200.77 | 1.02 | 9.90 |
| High TF expression, ATAC peak | 26.10 |  | 10.10 |  |
| Low TF expression, ATAC peak | 0.91 | 24.18 | 3.50 | 2.06 |
| High TF expression, no ATAC peak | 22.00 |  | 7.20 |  |
| Low TF expression, ATAC peak | 0.91 | 28.68 | 3.50 | 2.89 |
| High TF expression, ATAC peak | 26.10 |  | 10.10 |  |
| High TF expression, no ATAC peak | 22.00 | 1.19 | 7.20 | 1.40 |
| High TF expression, ATAC peak | 26.10 |  | 10.10 |  |

In testis, both the presence of open chromatin and high TF expression are associated with an expression boost, but every pairwise comparison shows a smaller magnitude difference than in ovary, indicating that trans regulation influences expression in ovary more than testis. While the presence of ATAC-seq peaks correlates with gene expression, TF expression is generally both neccesary and sufficient for gene expression in ovary. Most genes in testis have low but genuine expression (FPKM>1) without nearby detectable ATAC-seq peaks or high TF expression, indicating that leaky transcription may be commonplace in testis. The same category of genes in ovary has a median FPKM of 0.13, negligible by comparison.

## Discussion

Our results shape the contours of a model where gene age correlates with tissue-specific determinants of gene expression patterns. Genes are typically born under a simpler regulatory machinery (cis-regulation with fewer TF binding sites (Zhao et al., 2014)), sufficient to drive expression in the testis but not other tissues. As a gene ages, it will likely recruit more trans-acting TF partners, strengthen existing cis-acting TF binding sites (Tuğrul et al., 2015), or gain novel binding sites (Levran et al., 2020; Trizzino et al., 2017). In ovary, the presence of ATAC-seq peaks alone does not correlate with increased expression without help of trans-acting members of a gene’s network. The accumulation of TF partners boosts expression in other tissues more than testis, lowering the probability that a middle-aged gene will be testis-biased. Of course, many TF partners are repressive, meaning that their expression would be anti-correlated with that of their target gene. Despite this, older genes with larger TF networks are expressed across a greater variety of tissues, and with consistently higher expression levels than younger genes. Old genes likely continue to recruit more TF sites and relationships, and complex regulatory machinery such as enhancers or insulators. These features only marginally increase expression in the permissive transcriptional environment of the testis, but will substantially increase expression in other tissues, especially ovary. This trend will lead to common ovary-bias of older genes. The age-related complexity of a gene’s TF network may therefore drive functional recruitment of young genes to the testis, and old genes to the ovary.

DroID does not show whether a TF-gene relationship is one of an activator or repressor. For each gene, we use the same set of TF interactions across every tissue, so the activator/repressor balance should not bias the expression/upstream TF expression relationship between tissues. While our TF expression measure does not consider whether a TF is an activator or repressor, it is still quite predictive of expression. Indeed, the fact that this measure shows a fairly robust Pearson’s r with gene expression across tissues might indicate that most of these relationships are activation, consistent with Zhang and Zhou’s finding that genes accrue activating TFs and activating epigenetic marks concurrently as they age (Zhang and Zhou, 2019). Our TF expression metric relies on the assumption that most TFs are not an activator of a gene in one tissue and a repressor of the same gene in another tissue. It does not require the assumption that all TF-gene interactions are activation. While TF expression would be a poor method to predict gene expression, TF expression is a useful method to compare the relationship between trans regulation and gene expression between tissues.

It is also true that younger gene are often less studied compared to older genes. Fortunately, the DroID TF-gene interaction database shows ChiP-chip profiles of known TFs, giving us a whole-genome holistic comparison of confirmed TF-gene interactions without regard to the age of the target gene. This means that if a TF binds to the promoter of a younger gene, we will still be able to experimentally confirm this interaction even if the gene’s function is unknown. This high-throughput approach means that while we have a comprehensive list of confirmed TF-gene interactions, the activation/repression relationship and network modularity of many of these interactions is not yet known barring future lower-throughput experiments.

It has been proposed that the testis is uniquely positioned to drive the evolution of new genes due to an open chromatin environment (Assis, 2019; Kaessmann, 2010). Our findings indicate that this general pattern of open chromatin may be reflected on local levels, where we find a substantial proportions of genes of all age groups with ATAC-seq peaks in their promoters. Our findings indicate, however, that TF expression is more predictive of ovary gene expression than the presence of an ATAC-seq peak. This indicates that trans-regulation is especially important for ovarian gene expression, moreso than for testis expression.

Even though TF expression is higher in ovary than testis for every age group of genes, this activity does not result in ovary bias for young and middle-aged genes. A fitting analogy is that testis gene expression is like a bicycle in a low gear: easy to initiate movement, but total speed is limited despite the rider’s best efforts. Ovary gene expression is more like a bike in high gear: hard to initiate, but given a favorable environment (like biking downhill) the rider can reach greater speeds as a function of their energy input. This may explain why in long-term, ovary becomes a top niche for older genes.

Since a good number of the genes in this study originated before multicellular organisms (and therefore animal tissues such as testis and ovary), it is intriguing that such genes are affected by the relationship between gene age and tissue-specificity. Our results do not mean that the fate of all genes is to evolve in testis and gain expression in the ovary. Our results are a snapshot of the relationship between gene age and expression pattern as it occurs now, not a reconstruction of a guaranteed path for the evolution of a given gene’s expression.

It is instead clear that properties related to gene age differentially influence a gene’s potential roles in various tissues. Young genes have relatively few TF binding sites, a state not conducive to expression in most tissues except the testes. Older genes accumulate more TF binding sites (Tuğrul et al., 2015) and gain expression in non-testis tissues. Eventually, adding TF binding sites yields diminishing returns as a gene approaches expression saturation in a tissue. In ovary, however, added TF activity boosts expression more than in other tissues, making ovary-biased expression more likely for older genes. In testis, by comparison, adding TF binding sites appears to have a marginal effect on expression.

Future work could focus on the transcription factor aspect of this model. Given that old genes have more TFs than young genes, we would aim to simulate the evolution of a gene’s expression trajectory by adding a variable number of TF sites to the promoter of a reporter construct, and analyzing the tissue-specific expression patterns of the construct. This could tell us about the probable evolutionary “fate” of a stereotypical gene’s expression: to originate with testis-bias, gain expression in every other tissue, but end with highest expression in ovary. Why ovary currently becomes the top niche remains enigmatic and warrants future studies.

Of course, gene expression evolution takes place over millions or billions of years. Newly-originated genes, if they reach fixation in the population, will likely acquire TF sites over time. In another billion years, the regulatory characteristics that today confer testis-bias or ovary-bias may confer bias towards other tissues or even tissues that have not yet emerged.

## Methods

### Processing of FlyAtlas2 RNA-seq data

Fastq files of adult FlyAtlas2 tissues were obtained from EBI under accession number PRJEB22205 and reads were trimmed with Trimmomatic, set to remove the Illumina universal adapter. Reads were aligned with Hisat2 (Kim et al., 2016), default parameters to the Flybase dmel-r6.15 genome assembly (Thurmond et al., 2019). Reads with mapping quality less than 10 were removed. FPKM values were calculated with Stringtie (Kim et al., 2016) using default parameters. For each gene, FPKMs were averaged across replicates of a tissue.

### Determination of consensus gene ages

To allow for better statistical power and relatively uniform group sizes between gene age groups we binned genes into 3 groups: genes that emerged after the pan-Drosophilid divergence (Drosophilid), genes that emerged sometime before the pan-Drosophilid divergence but before the divergence of Bilateria (pre-Drosophilid), and genes that emerged before Bilateria (pre-Bilateria). To define Drosophilid genes, we used genes assigned to branches 1-5 in the gene age dataset from Zhang et al. (Zhang et al., 2010b). Ages of older genes were assigned using gene ages from Kondo et al (Kondo et al., 2017). Genes without ages defined in either dataset were not included for figures that segment genes by age, but were included for analyses of TF expression and open chromatin that did not consider gene age. For supplemental figures we reproduced the main figures defining genes from all 3 age groups only according to the ages assigned by Kondo et al. (Kondo et al., 2017) and observed no differences that would change our main findings.

### Calculation of tissue specificity

We used the tau method (Kryuchkova-Mostacci and Robinson-Rechavi, 2017) to calculate tissue specificity based on a gene’s FPKM across adult tissues, with replicates averaged (Kryuchkova-Mostacci and Robinson-Rechavi, 2017). A tau close to 1 indicates a tissue-specific gene, with a tau of 1 indicating a gene is only expressed in one tissue. A tau close to zero indicates a gene is equally expressed in every tissue.

### Calculation of scaled gene expression

FPKM is not normalized between genes, so we scaled gene expression to compare genes with different thresholds of activity. For a tissue i, scaled expression of a gene j is log-transformed FPKM in tissue i divided by gene j’s max logFPKM in any tissue. A scaled expression of 1 is gene j’s maximum expression in any tissue, and a scaled expression of 0 means expression is not detected. A scaled expression of 0.5 means that the logFPKM of a gene in a particular tissue is half the maximum observed logFPKM in any tissue.

### Calculation of TF expression for genes/tissues

For each gene, we wanted a measure for the activity of its upstream regulators in every tissue. We used the DroID database (Murali et al., 2011), which, for over 700 TFs, lists all genes whose promoters are bound by each TF as annotated with ChIP-chip and ChIP-seq by the modENCODE project (Roy et al., 2010). For this analysis, we only used genes with at least one TF annotated by DroID.

For a gene in a tissue, the TF expression score is the summed scaled expression of all annotated TF partners of that gene in that tissue. For example: if a gene’s TF partners have scaled expression values of 1, 1, and 0.5 in a tissue, and 0, 0, 0.5 in another tissue, the activity score for that gene would be 2.5 in the first tissue and 0.5 in the second, reflecting higher network activity in the first tissue. Since the TF expression values are scaled first, this measure allows for holistic comparisons of TF expression patterns between genes and tissues. The correlation between open promoter chromatin and TF expression in multiple tissues assures us that this metric measures biologically meaningful activity.

### ATAC-seq of Drosophila testis

We performed ATAC-seq experiment and analysis using 2-day-old testis of *D. melanogaster* RAL517 stain. For each sample, 25 newly emerged males were collected and transferred to 3 new vials (performed in triplicate). 48 hours later, we dissected testes in cold PBS. Tissues were lysed in 200 µl of ATAC-Seq lysis buffer (10mM Tris-HCl, 10mM NaCl, 3mM MgCl2, 0.1% IGEPAL CA-630) and manually homogenized with a plastic pestle, followed by a 1-minute incubation on ice, this process was repeated three times. The samples were pelleted at 4°C (100g for 10 minutes) to recover the nuclei. The buffer was removed and the nuclear pellet was re-suspended in 200 µl of lysis buffer. The nuclei preparation was filtered through a 30 µm Nitex nylon mesh (Genesee Scientific #57-105); the filter was further washed with another 200 µl of lysis buffer to ensure optimal nuclear recovery. The purified nuclei were isolated by centrifugation at 1000g for 10 minutes at 4°C). Following buffer removal, the nuclei were processed for the tagmentation reaction by adding: 12.5 µl Nextera Tagment DNA Buffer, 11.25 µl ddH2O and 1.25 µl Tn5 Transposase (Illumina Kit # FC-121-1030). The reaction was carried out in a thermal cycler for 30 minutes at 37°C with an additional mixing step 15 minutes into the reaction. The fragments were then purified using the Qiagen MinElute PCR purification kit (#28004) according to instructions. Libraries were constructed using the same primers as Buenrostro et al. (Buenrostro et al., 2015) and following a similar workflow: the purified DNA was first amplified for 5 cycles by PCR using the NEB Ultra II PCR mix (M0544). Then, an aliquot of the PCR reaction was analyzed by qPCR to determine the remaining optimal number of PCR cycles. Libraries were finally purified using SPRI beads with a two-step size selection protocol with bead-to-sample ratios of 0.55× and 1.00× for the first and second step, respectively. An aliquot of the purified library was used for quality control, and tested on an Agilent D1000 Tapestation platform, where concentration and peak periodicity were assessed. The samples were additionally tested for quality using Qubit, and sequenced on a 75bp paired-end Hiseq X platform.

### Processing of ATAC-seq data from testis, ovary, and S2 cells

We generated 3 replicates of testis ATAC-seq data from *D. melanogaster*. 2 replicates of OSC data were used: SRR3503078 and SRR3503086 (Iwasaki et al., 2016). 2 replicates of S2 cells were used: SRR5985082 and SRR5985083 (Ibrahim et al., 2018). Reads were aligned with bowtie2 (Langmead and Salzberg, 2012), default parameters against the flybase dmel_r6.24 reference genome (Thurmond et al., 2019). BAM files for each tissue were then merged with samtools merge (Li et al., 2009). Macs2 (Zhang et al., 2008) was used to call peaks for each tissue with the –nomodel parameter. The narrowpeak files were then loaded into R for further processing with Chippeakanno (Zhu et al., 2010) (details in supplementary Rmd on Github). Only peaks with a q value < 0.05 were used. Chippeakanno was run to find peaks overlapping the region 2000 bp upstream – 100 bp downstream of every gene’s TSS.

## Data availability

Scripts and processed data needed to reproduce figures are deposited in https://github.com/LiZhaoLab/TissueSpecificity. Testis ATAC-seq of *Drosophila melanogaster* Ral517 is deposited at NCBI under biosample accession # SAMN16259271.

## Data reproducibility

The data needed to reproduce this work can be found in this link https://github.com/LiZhaoLab/TissueSpecificity. It includes calculated FPKM for every gene and tissue in FlyAtlas2, files used to calculate consensus gene ages from Kondo et al. and Zhang et al., narrowpeak files we calculated for each of the 3 ATAC-seq datasets, a csv file with calculated TF expression (connectivity.csv) for each gene and tissue, and a file from DroID showing every experimentally annotated TF-gene interaction (tf_gene.txt). These files are all referenced by the Rmd script on our Github page. The free web app which allows users to interactively explore our tissue specificity data for any set of genes without coding experience necessary is: https://zhao.labapps.rockefeller.edu/tissue-specificity/.

## Acknowledgements

We are grateful to Stuart McDonald from The University of Kansas and Margo Herre from Leslie Vosshall’s lab at The Rockefeller University for the help and suggestions on ATAC-seq, and Luis Gracia at The Rockefeller University for the help with the web app. We thank the members of the Zhao lab for helpful discussions and critically reading an earlier version of the manuscript.

## Author contribution

E.W. and L.Z. conceived the study. N.S. and S.B. generated the ATAC-seq data. E.W. performed all the analysis in this manuscript. E.W. and L.Z. wrote the manuscript with the input from all authors.

## Supplementary figures and tables for

**Supplemental Figure 1:**
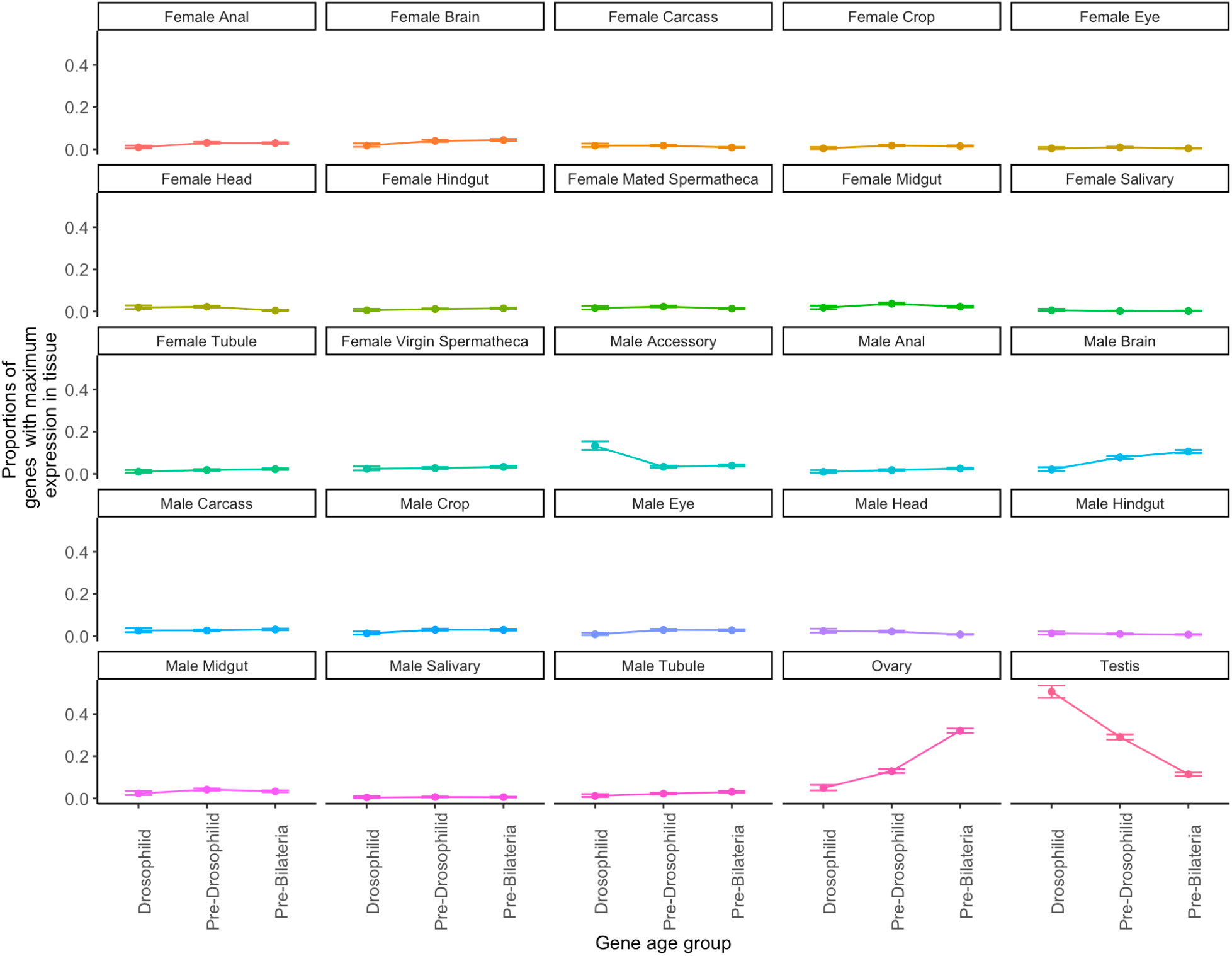
Proportion of genes with maximum expression in a tissue, by age group. A high proportion of young genes have maximum expression in testis, and a high proportion of older genes have maximum expression in the ovary. Between the testis and accessory gland, a majority of Drosophilid genes are biased towards the male reproductive system. Some tissues like male brain and male accessory gland show a trend in age-related tissue bias, but none as large as the testis and ovary.

**Supplemental Figure 2:**
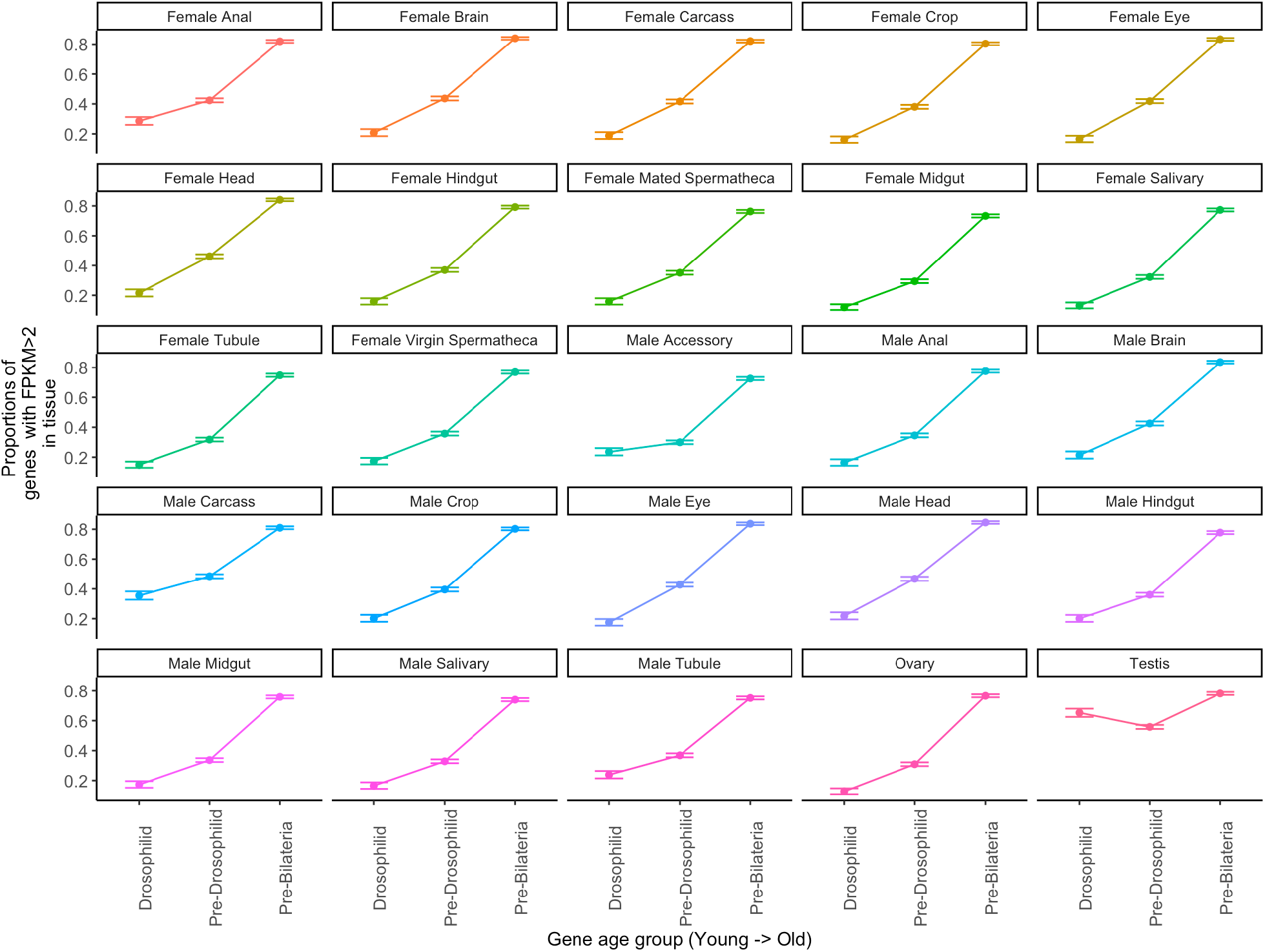
Proportion of genes expressed with FPKM > 2, by Fylatlas2 tissue and age. In every tissue, old genes are most commonly expressed, but ovary has the greatest difference between young and old genes, and testis has the smallest difference.

**Supplemental Figure 3:**
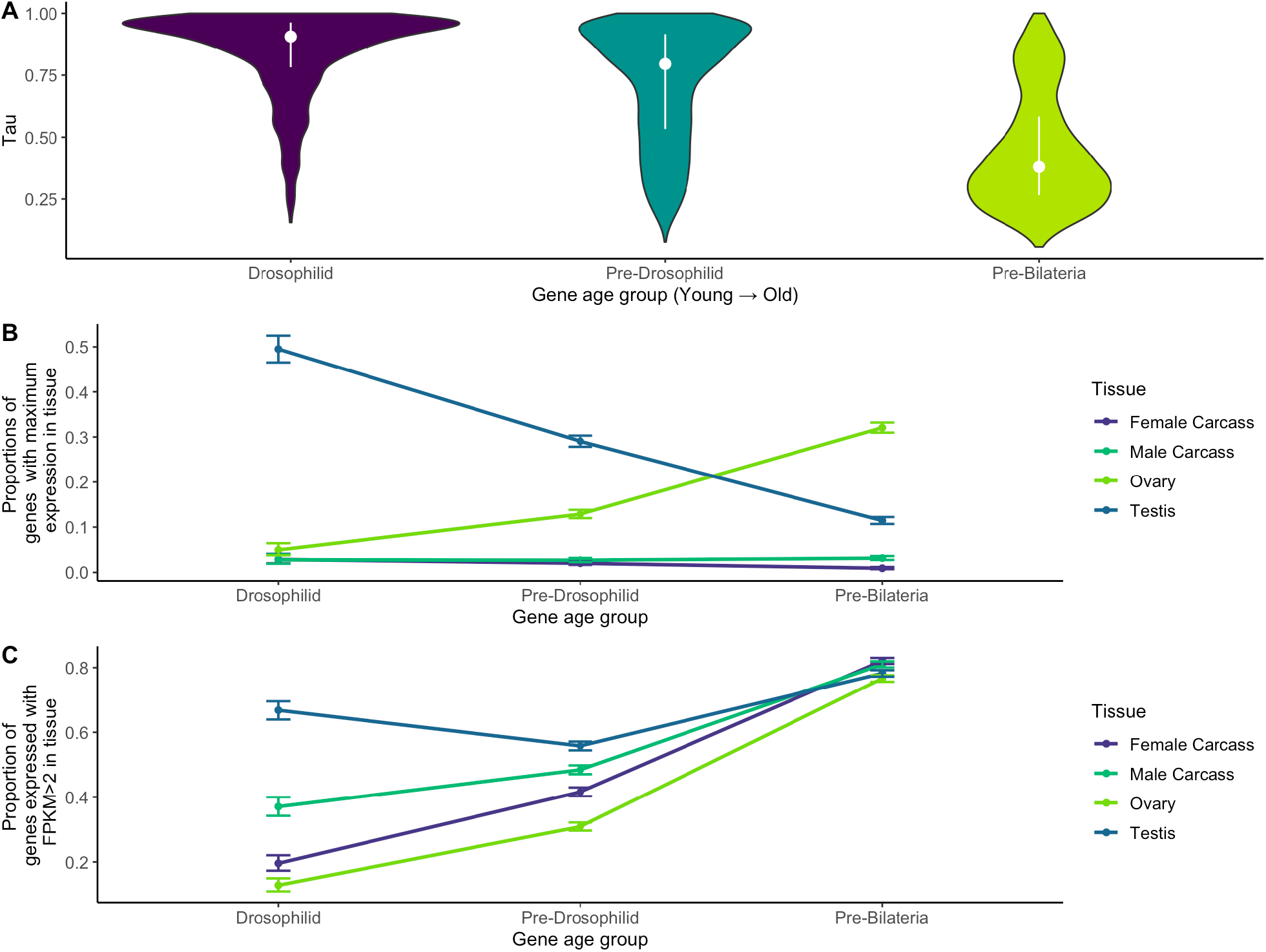
Young duplicate genes do not confound results from Figure 1. Recently duplicated genes may have high sequence similarity to their parent copies, causing mapping ambiguities. Shown is the analysis from Figure 1, with genes annotated by Kondo et al. as “*melanogaster*-only” removed. No conclusions from Figure 1 are changed, indicating that *melanogaster-*specific genes do not confound the high testis-specificity of Drosophilid genes.

**Supplemental Figure 4:**
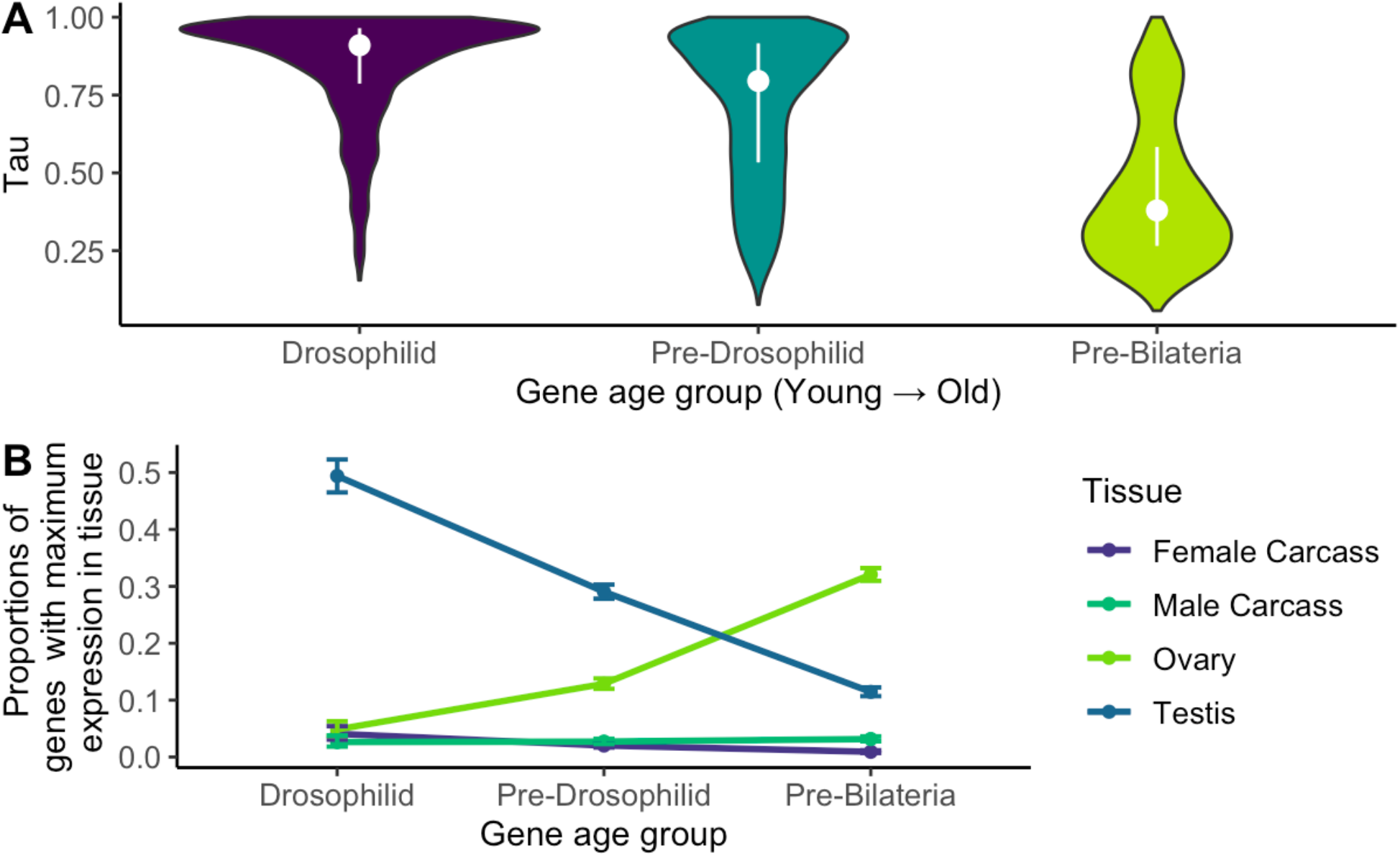
Alternate gene age assignments confirm patterns of tissue bias and tissue specificity. The main figures assign Drosophilid genes as those characterized by Zhang et al. and use ages from Kondo et al. for older genes. We remade Figures 1A and 1B with all gene ages assigned from Kondo et al. and found that young genes are more tissue-specific than old genes, young genes are often testis-biased and old genes are ovary biased.

**Supplemental Figure 5:**
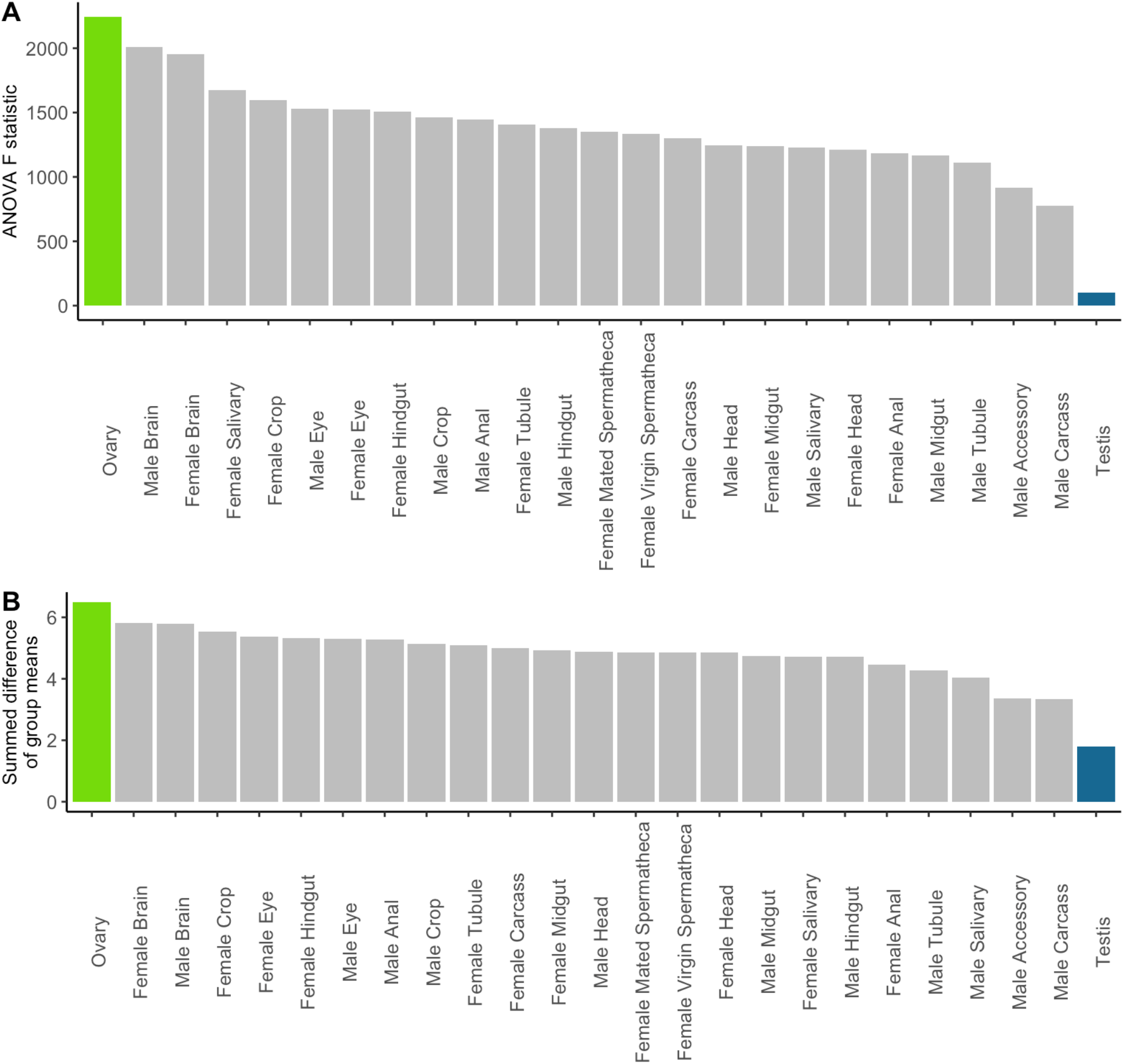
Very young genes do not confound results from Figure 2b. With *melanogaster*-specific genes removed, gene expression varies the least between age groups in testis, and most in ovary, just as in the main text. A.) ANOVA F statistic between tissues. B.) Summed difference of mean expression between age groups in each tissue.

**Supplemental Figure 6:**
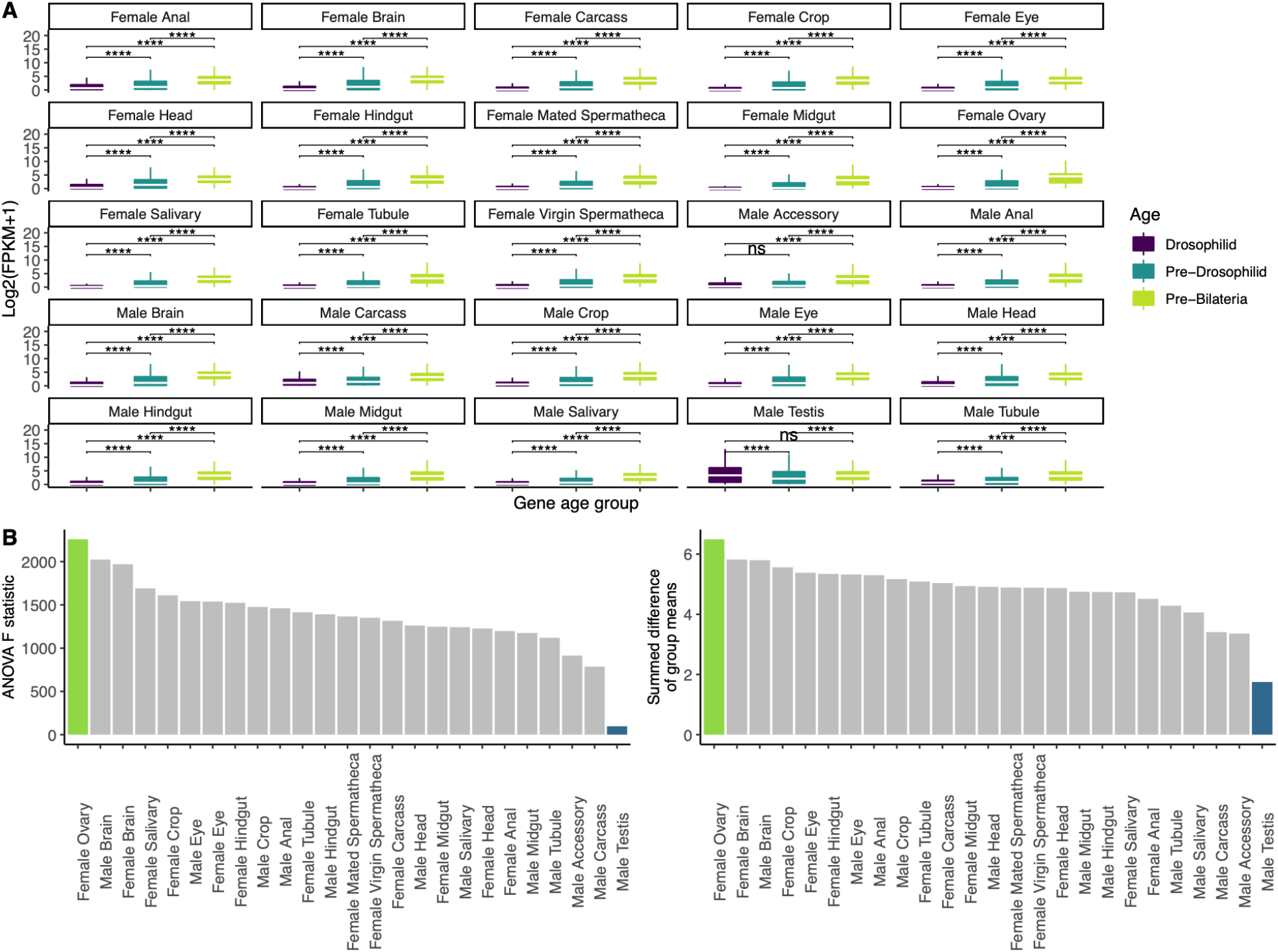
Alternate gene age assignments do not alter main conclusions from Figure 2. This is the analysis from Figure 2 using only gene age assignments calculated by Kondo et al. In part A, two things have changed. Drosophilid genes have statistically similar expression to Pre-Drosophilid genes in accessory glands, the other male reproductive tissue. In testis, Drosophilid genes are statistically similar to Pre-Bilateria genes instead of Pre-Drosophilid genes. Neither of these changes affect the results from part B, which, like the main figures, shows that ovarian gene expression varies more with age than any other tissue, and testis gene expression varies the least.

**Supplemental Figure 7:**
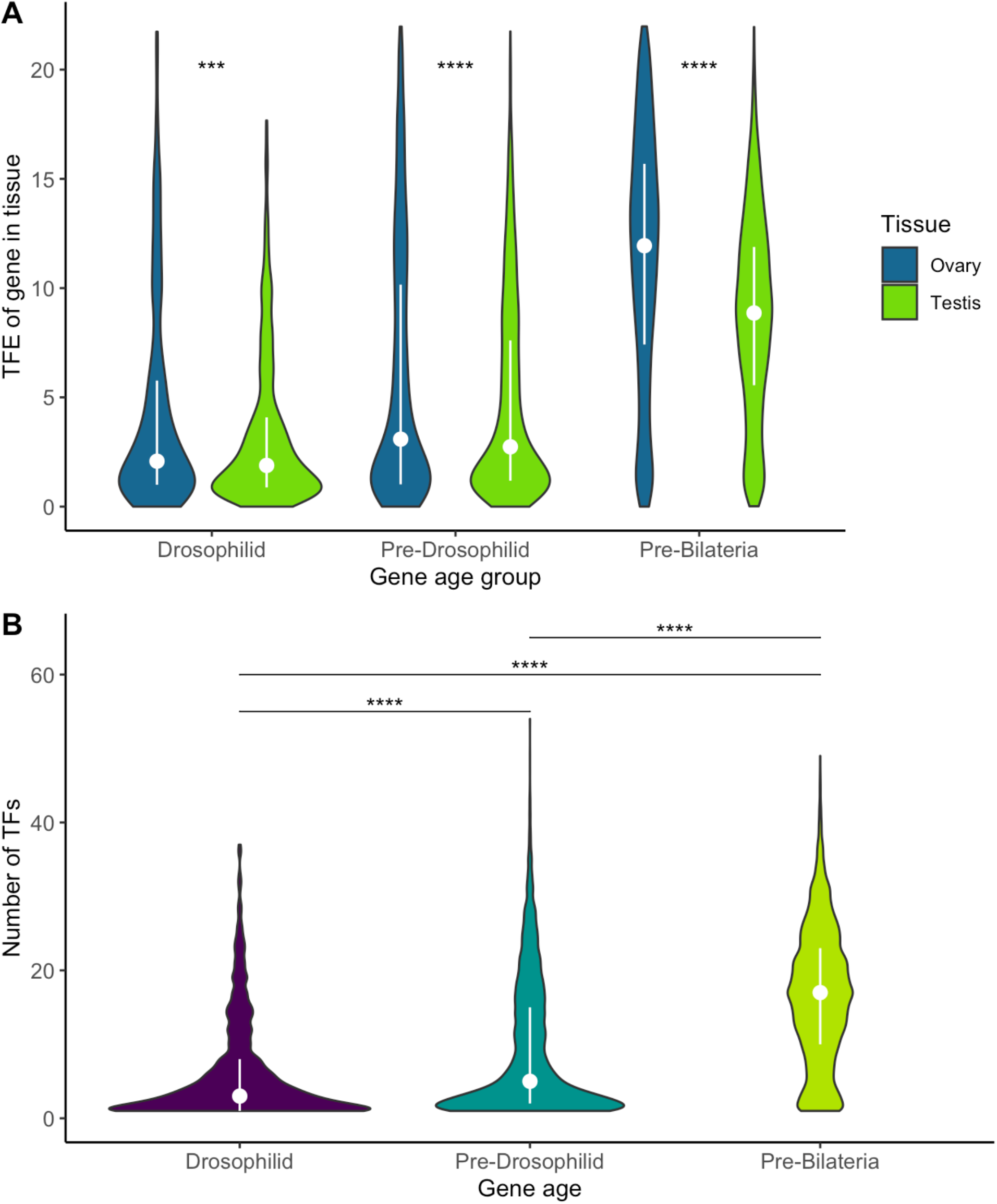
Alternate gene age assignments do not affect results from Figure 3A and 3B. Using gene ages calculated by Kondo et al., we found that no age group of genes shows elevated TF expression in testis compared to ovary (A). Additionally, the promoters of young genes are less likely to be bound by known TFs than older genes (B).

**Supplemental Figure 8:**
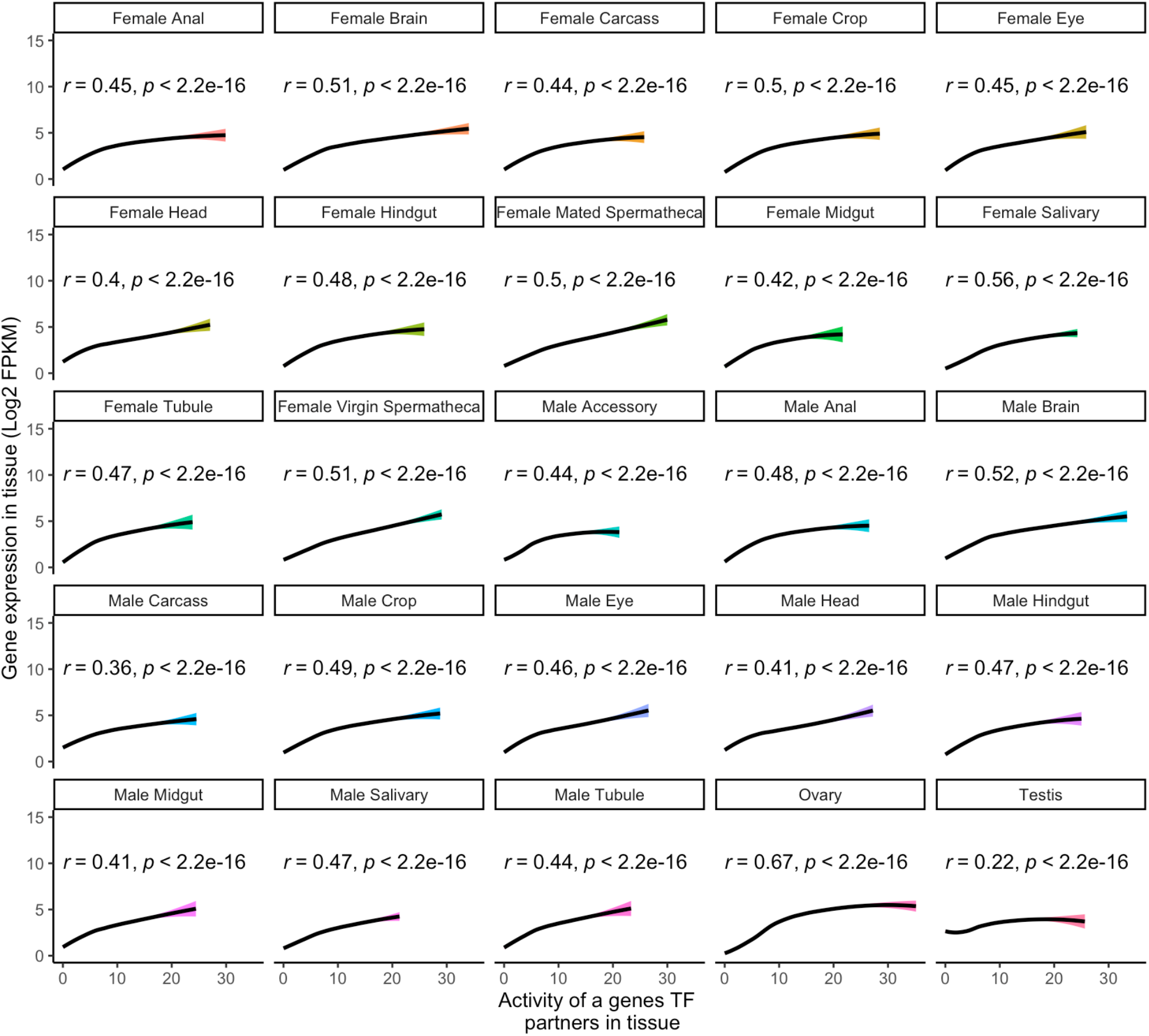
TF expression vs gene expression in every Flyatlas2 tissue. Corresponding to Figure 3, this figure compares TF expression to gene expression across all tissues. Gene expression is least responsive to TF expression in testis, and most responsive to TF expression in ovary. Ovary also has the largest range of TF expression. Lines are a smoothed loess regression with 95 percent confidence intervals. Pearson’s r is shown for every tissue, and it is largest in ovary and smallest in testis.

**Supplemental table 1:**
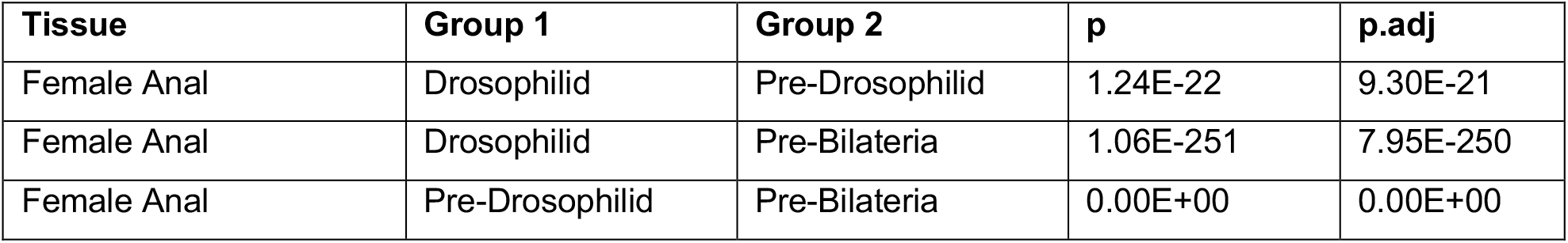
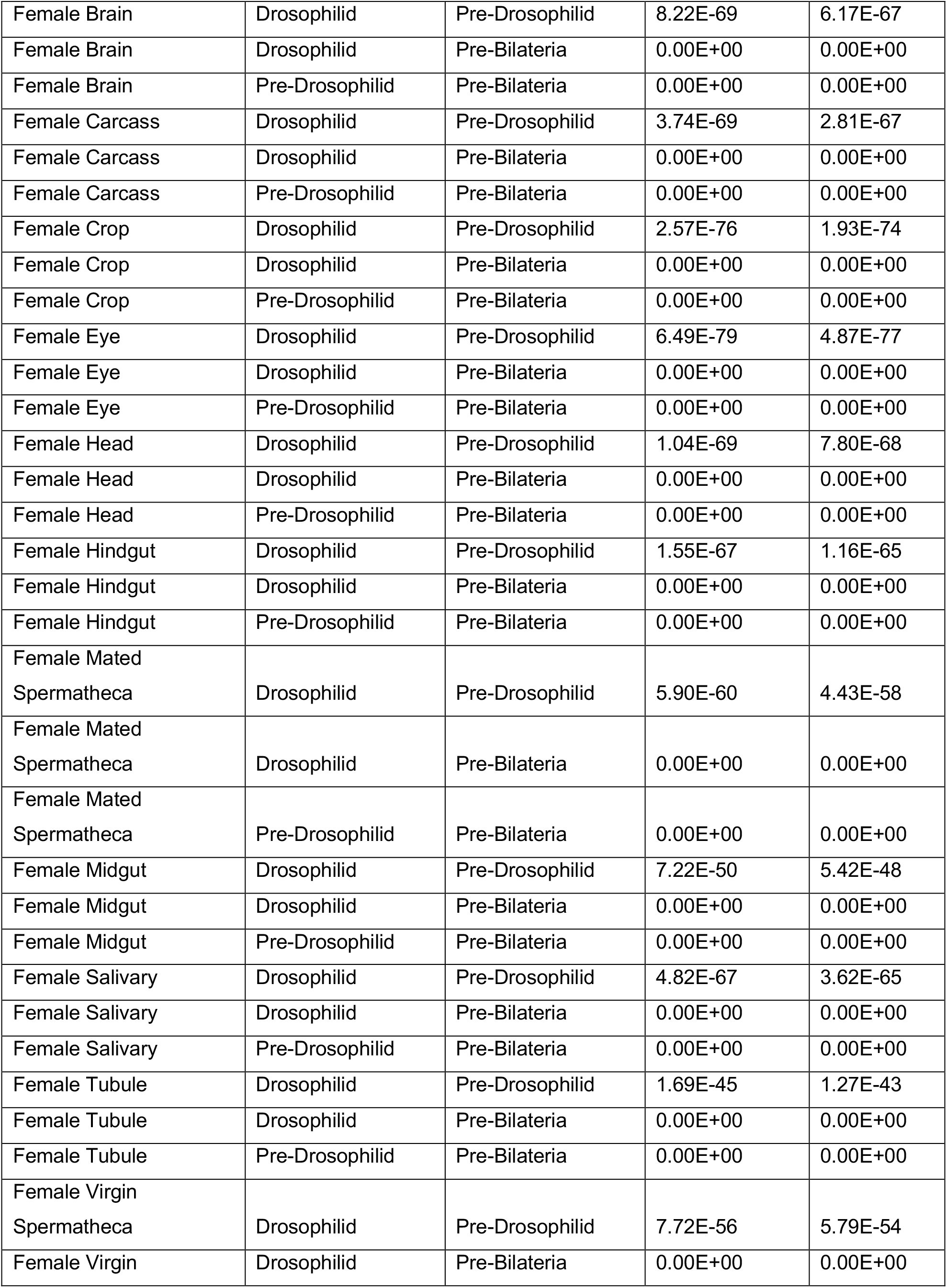
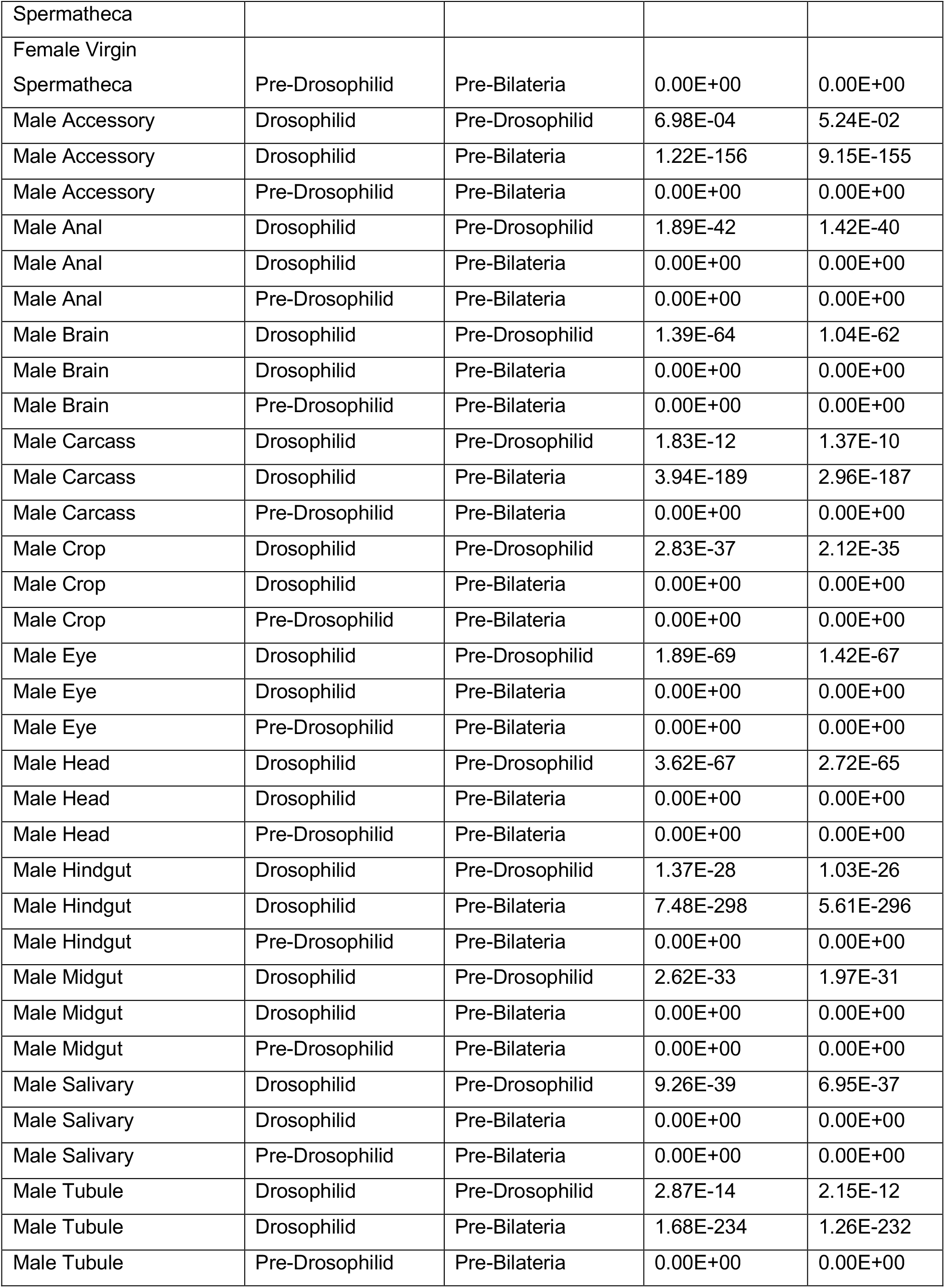
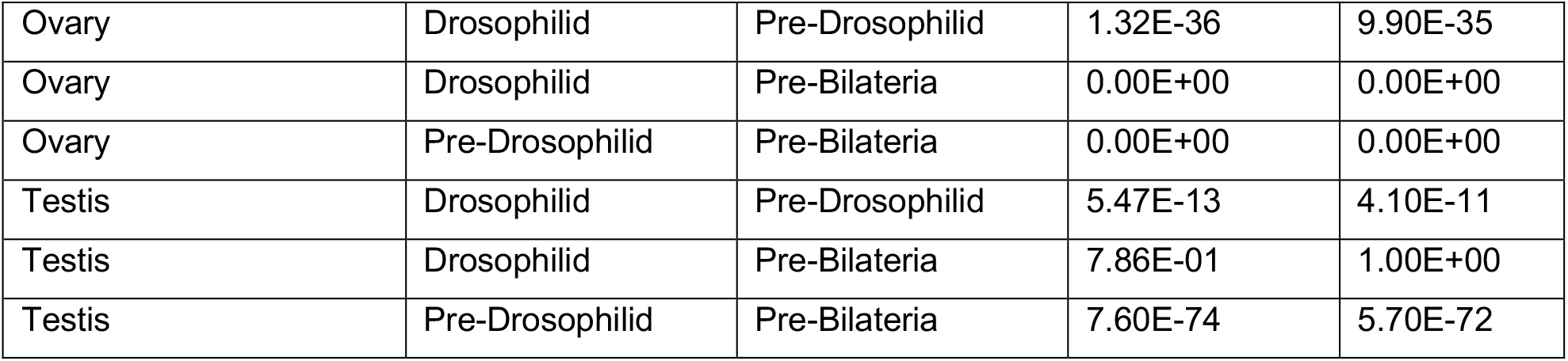
Raw and adjusted p values for Figure 2. For each tissue and pairwise comparison shown in Figure 2. These are the raw and Bonferroni-corrected p values for each comparison.

